# Integrating Fungal–Bacterial Synergy to Enhance Circular MFC–Hydroponic Performance

**DOI:** 10.64898/2026.03.23.713689

**Authors:** Ignacio Baquedano, Daniel González-García, Alicia Prieto, Jorge Barriuso

## Abstract

Microbial fuel cells (MFCs) represent a promising technology for the simultaneous treatment of wastewater and bioelectricity generation. In this study, the MFCs are conceived as functional modules to be integrated into hydroponic cultivation systems, acting as a prosthetic rhizosphere capable of coupling wastewater treatment and bioelectrochemical activity with plant nutrition improvement.

We compared the electrochemical performance of different microbial consortia comprising the electroactive bacterium *Shewanella oneidensis,* the plant growth promoting rhizobacterium (PGPR) *Pseudomonas putida,* and the plant biomass-degrading fungus *Ophiostoma piceae*, along with the supplementation with the quorum sensing (QS) analogue molecule 1□ dodecanol. These microbial consortia are tested in MFCs fed with wastewater and root exudates to analyze enhanced feedstock assimilation, electricity production, and the generation of plant growth–promoting substances (PGPS).

From an electrochemical perspective, we evaluated planktonic growth, anode adhesion, substrate consumption, and the production of redox-active molecules and PGPS such as flavins and siderophores respectively alongside key electrical production parameters, including current output and power. Among the different microbial configurations tested, the consortium combining *S. oneidensis*, *P. putida*, and *O. piceae* exhibited the highest electrical production potential. Moreover, within this framework, we detected the extracellular production of siderophores in MFCs containing *P. putida*, suggesting a potential role supporting hydroponic crop growth. Furthermore, the addition of 1-dodecanol led to an improvement of the bioelectrochemical parameters.

These results highlight the potential of synthetic microbial consortia in MFC-based systems not only to enhance electricity generation from wastewater but also to provide added value in integrated hydroponic applications through rhizosphere-like functions.

## INTRODUCTION

Microbial fuel cells (MFCs) represent a promising bioelectrochemical technology capable of addressing several environmental and energetic challenges simultaneously. By exploiting the metabolic activity of electroactive microorganisms, MFCs enable the direct conversion of chemical energy stored in organic matter into electrical energy (Bond et al., 2002; Logan, 2009; Lovley, 2006).

The performance of MFCs strongly depends on the substrate feeding the anode and on the structure of the anodic microbial community. Electricity generation relies on extracellular electron transfer (EET) processes, whereby microorganisms attached to the anode transfer electrons either directly to the anode surface by direct electron transfer (DET) or indirectly by planktonic microorganisms via redox mediators (Kumar et al., 2017; Shi et al., 2016). In addition to microbe–electrode interactions, electron transfer can also occur between different members of the microbial community through metabolic cooperation, redox shuttles, syntrophic interactions or Direct Interspecies Electron Transfer (DIET), where electrons are exchanged via conductive pili or extracellular nanowires without the need for soluble mediators (Lovley, 2011, 2017). These complex interactions highlight the importance of community composition and metabolic diversity in determining both the electrochemical output and operational stability.

MFCs have been explored for wastewater treatment and for the recovery and production of value-added compounds, thereby improving the sustainability and economic viability of this technology. Urban wastewater represents one of the most abundant and complex anthropogenic waste streams, containing high loads of organic matter, nutrients, and diverse chemical contaminants (Sevda et al., 2013; Yuan et al., 2016). Agricultural wastewaters are also of particular concern, as they often contain elevated concentrations of nitrates and phosphates derived from fertilizers, contributing to eutrophication and groundwater contamination (Camargo & Alonso, 2006; Carpenter et al., 1998). Effective treatment of these effluents is essential to protect aquatic ecosystems and public health, while also aligning with circular economy strategies that promote resource recovery rather than simple disposal. In this context, MFCs complement conventional wastewater treatment technologies.

Recently, the integration of MFCs into hydroponic cultivation systems has been proposed as a novel strategy to couple wastewater treatment, energy generation, and soilless agriculture (Paucar & Sato, 2022; Sato et al., 2023). Within such systems, the anodic chamber of MFCs can function as a prosthetic rhizosphere, mimicking ecological and biochemical functions of the natural root–soil interface. These include the establishment of beneficial microbial communities that contribute to nutrient mobilization, plant elicitation through signaling molecules, and disease suppression mediated by niche occupation and antagonistic activity against plant pathogens. This artificial rhizosphere can host metabolically active microbial communities capable of degrading organic compounds present in wastewater while producing plant growth–promoting substances (PGPS), including phytohormones, amino acids, and microbial metabolites that enhance nutrient uptake, stress tolerance, photosynthesis, and overall plant fitness (Paucar & Sato, 2022). This can be done either directly or through PGPR-mediated functions such as nitrogen fixation and phosphorus solubilization. Among these, siderophores—low-molecular-weight iron-chelating compounds—play a key role in enhancing Fe^3+^ bioavailability and their positive effects in hydroponic systems are well known (Crowley et al., 1992; Stegelmeier et al., 2022).

In this context, the system can operate as a closed loop in which electricity produced by the MFCs powers LED lighting for plant growth, while a continuous recirculation flow connects the MFC anode with the hydroponic medium. This configuration enables bidirectional exchange: beneficial metabolites generated by the anodic microbial community are delivered to the plants, and root exudates are subsequently returned to the anode, where they sustain microbial activity and electron transfer. Such a loop fosters synergistic interactions that enhance both energy recovery and crop fitness. The integration of MFCs into multifunctional platforms therefore creates opportunities to simultaneously generate electricity, remediate wastewater, and improve plant performance through microbial metabolic processes (Fig. 1).

**Figure 1.**
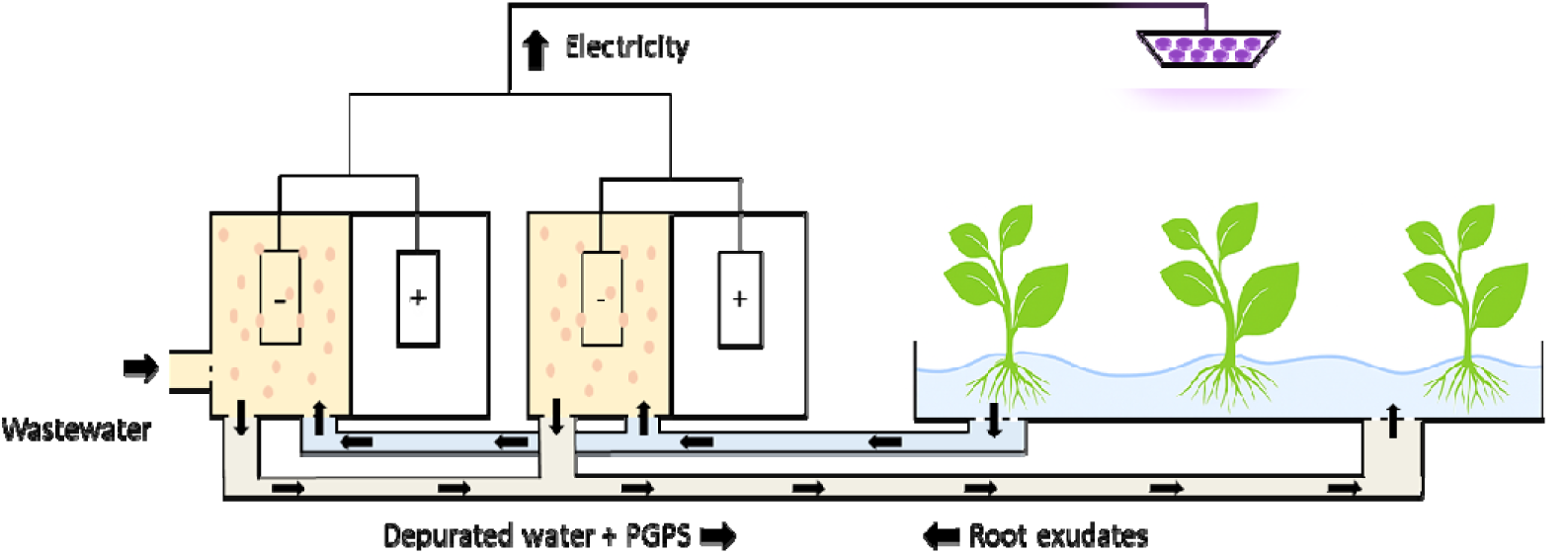
Schematic representation of the integration of MFCs into hydroponic cultivation systems, in which wastewater, hydroponic medium and root exudates feed the anode chamber of MFCs. Within the anode, the microbial community functions as a prosthetic rhizosphere that performs three interconnected functions: (i) wastewater depuration; (ii) bioelectricity generation; and (iii) production of plant-beneficial compounds.

From an ecological and functional perspective, microbial consortia offer advantages over single strains. Structured communities enable the distribution and compartmentalization of metabolic tasks among members with complementary capabilities, improving substrate utilization, degradation of complex compounds, PGPS production, and biofilm stability.

These properties are particularly relevant for wastewater-fed MFCs, where substrate complexity and variability are high (Getu et al., 2025; Modesto et al., 2026).

Filamentous fungi represent promising yet underexplored components of anodic consortia. Their extensive enzymatic repertoires allow the degradation of lignocellulosic and recalcitrant substrates, releasing simpler metabolites that can fuel electroactive bacteria. Although bacteria dominate most MFC studies, several fungi have demonstrated electrochemical activity or have been successfully implemented in MFC systems. Species such as *Saccharomyces cerevisiae*, *Candida spp*., and *Yarrowia lipolytica* have been reported to participate in electron transfer, typically mediated by endogenous or exogenous redox shuttles. Some filamentous fungi, including white-rot species, have also shown potential for enhancing electron transfer through the production of redox-active metabolites. However, filamentous dimorphic fungi remain largely unexplored in MFC configurations, particularly in multispecies consortia (Sekrecka-Belniak & Toczyłowska-Mamińska, 2018). In this study, we focus on three microorganisms with complementary metabolic and functional traits: i) *Ophiostoma piceae*, an aerobic wood-saprophytic dimorphic fungus with a broad enzymatic arsenal, which uses quorum sensing (QS) regulation to control the morphological switch and the secretion of enzymes (De Salas et al., 2015), ii) *Pseudomonas putida* KT2440, an aerobic metabolically versatile bacterium that effectively colonizes the rhizosphere and exhibits PGPR traits, including siderophore production (Costa-Gutierrez et al., 2022). It also communicates via QS mechanisms, participates in electron transfer through mediators and DIET, and forms stable biofilms. Furthermore, it has been demonstrated that this bacterium is able to associate with *O. piceae* (Ruiz et al., 2021), iii) *Shewanella oneidensis*, an extensively used model electroactive bacterium due to its efficient EET mechanisms, which occur either directly through outer-membrane c-type cytochromes and conductive nanowires —facilitated by its ability to adhere to surfaces— or indirectly via soluble electron shuttles such as riboflavins (Marsili et al., 2008; Shi et al., 2012). This facultative anaerobe exhibits a comparatively restricted metabolic versatility relative to previously described species. Besides, regulation of population density has been described by QS-mediated regulation. The objective of this work was to achieve efficient wastewater treatment while simultaneously promoting electricity generation and production of metabolites of interest for hydroponic crops. Combinations of the three different microorganisms were tested in consortia, and the effect of adding 1-dodecanol, an analogue of the quorum sensing molecule (QSM) dodecanoic acid, was evaluated. The performance of the consortia cultivated in an artificial medium was assessed in terms of substrate metabolization, growth, biofilm formation, electrical output, and riboflavin and siderophores production. By comparing single- and multi-species systems in the presence or absence of a QS molecule, we aim to elucidate the potential benefits of engineered microbial communities for integrated MFC-hydroponic applications.

## MATERIALS AND METHODS

### Pre-inoculum growth conditions

*O. piceae* was routinely grown in YPD medium composed of yeast extract (10 g·L□¹), peptone (20 g·L□¹), and dextrose (20 g·L□¹). *P. putida* KT2440 was cultivated in LB medium containing tryptone (10 g·L□¹), yeast extract (5 g·L□¹), and sodium chloride (10 g·L□¹). *S. oneidensis* MR-1 was grown in TSB medium consisting of casein peptone (pancreatic digest) (17 g·L□¹), soya peptone (papain digest) (3 g·L□¹), glucose (2.5 g·L□¹), dipotassium hydrogen phosphate (2.5 g·L□¹), and sodium chloride (5 g·L□¹).

All cultures were incubated at 28 °C with orbital shaking at 180 rpm to the appropriate growth and used as inoculum for the experimental assays.

### Design of the anodic medium

The anodic medium was formulated to simulate a wastewater-fed hydroponic system and consisted of 45% artificial urine (Brooks & Keevil, 1997), mimicking urban wastewater, 45% Sonneveld hydroponic nutrient solution (Sonneveld et al., 1999), and 10% root-derived compounds and sloughing materials. The artificial urine was slightly modified by reducing phosphate and CaCl□ concentrations and adding sodium nitrate, while the Sonneveld solution was adjusted to lower nitrate concentrations. To simulate root-derived inputs, encompassing both soluble exudates and structural materials originating from root sloughing and cell wall turnover, cellulose, pectin, and hemicellulose were included as representative carbohydrates (Xiong et al., 2009). Ferulic acid, pipecolic acid, allantoin, and histamine were incorporated as functional compounds linked to plant–microbe interactions (McLaughlin et al., 2023; Ren et al., 2016; Sun et al., 2022; Wang et al., 2021). The detailed composition of the medium is provided in Table S1.

### Planktonic growth and biofilm formation assays

Prior to experiments in the MFCs, planktonic growth and biofilm-forming capacity of the three microorganisms, individually and in consortia, was assayed using a standard 24-well microtiter plates system. One mL of the medium designed for the anode was used per well. Microbial pre-cultures were prepared, washed in the anodic medium and finally inoculated to an initial optical density (OD□□□) of 0.05 (0.1 for *O. piceae*). Plates were incubated statically at 28 °C for 28 h (52h for *O. piceae*). After incubation, planktonic growth was quantified by measuring the OD□□□ of the culture supernatant with a UV-1900i Spectrophotometer (Shimadzu). Biofilm formation was assessed using a crystal violet staining assay. Briefly, after removal of the culture supernatant, wells were gently washed with distilled water to remove non-adherent cells. Biomass attached to the plastic was then stained with 1 mL of 0.1% (w/v) crystal violet solution for 15 min under constant agitation. To remove the non-retained dye wells were washed twice with distilled water. The retained dye was subsequently solubilized by incubation with 1 mL of absolute ethanol for 20 min under constant agitation. Biofilm biomass was quantified by measuring the absorbance of the resulting solution at 590 nm. Values were corrected by subtracting the OD_590_ of the corresponding culture medium. Data shown in the graphs represent the mean values of the biological replicates and error bars correspond to the standard. All measurements were normalized to microorganism□free control conditions.

In addition, to evaluate microbial adhesion and biofilm formation on the electrode material, pieces of carbon veil with a surface area of 0.25 cm² were placed into selected wells containing the anodic medium, incubated as described above, and processed as described in “Microscopic assessment of microbial interactions on the anode” section.

### Microbial fuel cell setup and operation

Dual-chamber H-type MFCs were used in this study. Each chamber (anode and cathode) had a working volume of 120 mL and was separated by a cation exchange membrane (Nafion® N-117, DuPont). Both anode and cathode electrodes were made of carbon veil, with a surface area of 15 cm² for the anode and 30 cm² for the cathode. Electrical connections were established using nickel–chromium (Ni–Cr) wire with a diameter of 0.51 mm. The catholyte consisted in a solution of 50 mM potassium hexacyanoferrate(III), 50 mM K□HPO□, and 50 mM KH□PO□. The external circuit was closed using a fixed external resistance of 2.7 kΩ. The anodic chamber was inoculated with 120 mL of a preculture of each microorganism, previously washed with anodic medium and adjusted to an initial OD□□□ of 0.5 before inoculation. This increase in the initial microorganism concentration was intended for a better bioelectrochemical performance given the short experimental time. When used, 1-dodecanol (Merck) was added at a final concentration of 50 µM. MFCs were operated at a constant temperature of 28 °C. Voltage output was monitored every 30 min using an ADC-24 data logger (Pico Technology, UK), while polarization curves were obtained using a NEV4 potentiostat (NanoElectra) by sweeping the cell voltage from 800 mV to 0 mV with a slope of −1 mV.

All experiments were conducted in two independent experimental rounds. In each round, one biological replicate of every MFC condition was performed. The values shown in the text correspond to the average value and the standard error.

### Microbial growth dynamics in MFC anolytes

Microbial growth in the anolyte of the MFCs was evaluated at the end of the experimental operation. Planktonic biomass was first estimated by measuring OD□□□ of the anolyte. The data correspond to the mean OD□□□ value and the standard error of the biological replicates.

### Microscopic assessment of microbial interactions on the anode

To visualize microbial colonization and biofilm architecture on the anode material, scanning electron microscopy (SEM) and confocal laser scanning microscopy (CLSM) analyses were performed. In both cases, carbon veil fragments (approximately 0.5 cm²) were collected from the anode at the end of the experiment. Immediately after excision, samples were washed twice with PBS to remove loosely attached cells and medium residues. For SEM analysis, the samples were fixed in 2.5% (v/v) glutaraldehyde overnight at 4 °C. After fixation, samples were washed twice with distilled water and subsequently dehydrated through 10 min incubation in growing ethanol concentrations (30, 50, 70, 80, 90, and 100%). Finally, the filters were subjected to critical point drying and metallization at the Spanish National Centre for Electron Microscopy (Universidad Complutense de Madrid) and images were acquired using a JEOL 6400 scanning electron microscope. For CLSM analysis, following the initial PBS washing, samples were fixed with 4% (w/v) paraformaldehyde for 1 h at room temperature. Samples were then washed twice with distilled water and incubated in the dark with propidium iodide (25 µg·mL□¹) for 30 min at room temperature. After staining, samples were washed twice with distilled water. Confocal images were acquired using a Leica TCS SP5 confocal microscope, exciting the fluorophore with a DPSS 561 laser and collecting emission signals between 577 and 655 nm. Images were obtained using a 20× objective, acquiring a total of 51 optical z-planes. The images presented correspond to maximum projections.

### Urea quantification by High-Performance Liquid Chromatography (HPLC)

At the end of the experiment, 500 µL of anolyte were collected and centrifuged to remove suspended biomass. The resulting supernatant was subsequently filtered prior to analysis.

HPLC analyses were performed using an Agilent 1200 Series system equipped with a refractive index detector (RID). 10□ µL of sample were injected and analyzed isocratically (0.5 mL/min) in a ZORBAX Eclipse Plus C18 column, with distilled water as the mobile phase. Urea was detected by RID with a retention time of approximately 5.7 min under the applied chromatographic conditions.

### Metabolite Analysis by Gas Chromatography–Mass Spectrometry (GC–MS)

Carbohydrates and other soluble metabolites contained in the anolyte were analyzed by gas chromatography–mass spectrometry (GC–MS). For each sample, 1.5 mL of anolyte were collected at the end of the experimental time, centrifuged to remove suspended biomass, and the supernatant was filtered. The filtrates were then completely dried in a SpeedVac and the samples were derivatized prior to analysis. Derivatization was performed with 250 µL of hydroxylamine hydrochloride in pyridine and incubated for 30 min at 70 °C. Subsequently, 250 µL of *N,O*-bis(trimethylsilyl)trifluoroacetamide (BSTFA) were added, and samples were incubated for 30 min at 80 °C. Glucuronic acid (150 ng) was used as an internal standard to ensure analytical consistency and semi-quantitative comparison among samples. GC–MS analyses were carried out using an Agilent 7980A gas chromatograph coupled to a 5975C mass selective detector. The results shown correspond to the average of two technical replicates and were normalized to the internal standard. Error bars correspond to standard error.

### Detection of flavins and siderophores

Riboflavins and siderophores were detected by fluorescence-based analysis of anolyte samples. For both determinations, 130 µL of anolyte were centrifuged to remove suspended biomass, and 100 µL of the resulting supernatant were transferred to individual wells of a 96-well microplate. Each sample was analyzed in triplicate using a microplate reader (SpectraMax M2). Riboflavin fluorescence was measured using an excitation wavelength of 450 nm and collecting emission at 530 nm. Siderophore-associated fluorescence was measured by excitation at 400 nm and emission detection at 455 nm. To specifically assess the metal-chelation-dependent component of siderophore fluorescence, the supernatant of each sample was pre-incubated with 2□ mM EDTA for 20□ min at room temperature prior to measurement. Data shown correspond to the mean values and standard error of the triplicates.

## RESULTS

### Design of the anodic medium

The MFCs were designed to be integrated into urban wastewater treatment and hydroponic cultivation systems. Under this configuration, the anode chambers are primarily fed with urban wastewater, and mixed with hydroponic nutrient solution and plant-derived root exudates. Based on this expected operational scenario, the final composition of the medium used in the experiments consisted of 45% modified artificial urine (Brooks & Keevil, 1997), with lactate, citrate and urea as the main components, 45% modified Sonneveld’s hydroponic solution (Sonneveld et al., 1999), and 10% root-derived compounds, including root exudates and potential sloughing materials (Table S1).

Given the limited chemical characterization of root-derived components in hydroponic systems, representative compounds described in the literature as general constituents of plant root exudates were selected to prepare the medium, while low-molecular-weight compounds expected to occur at very low concentrations were excluded. Instead, a limited set of functionally relevant molecules likely to modulate microbial interactions and metabolism was tested (McLaughlin et al., 2023; Ren et al., 2016; Sun et al., 2022; Wang et al., 2021; Xiong et al., 2009). This simplified anodic medium was designed to emulate a coupled wastewater–hydroponic environment, providing a chemical framework intended to support microbial growth, redox activity, and interspecies interactions. It included assimilable carbon sources and bioavailable nitrogen compatible with the metabolic capabilities of the selected microorganisms, as well as molecules representative of root exudates that could act as biochemical cues influencing growth, signaling, and potential syntrophic interactions within the consortia.

### Microbial growth and biofilm formation

To determine whether the selected anodic medium constitutes a suitable substrate to support microbial growth and potentially promote cooperative or syntrophic interactions, the growth capacity and biofilm-forming ability of the three selected microorganisms were evaluated individually and in consortia of two or three members. To mimic the physicochemical conditions at the anode chamber of MFCs, the microorganisms were grown in the designed anodic medium under static conditions, to limit oxygen availability. In this experimental setup, *O. piceae* exhibited limited apparent growth, as reflected by low optical density values at both 600 nm (0.5) and 590 nm (0.8) (Fig. 2). Although these values would initially suggest poor growth, microscopic examination of planktonic cultures revealed that growth occurred predominantly in the hyphal form. In contrast, both bacterial species, *S. oneidensis* and *P. putida*, displayed more robust growth, with high planktonic densities (OD□□□ of 2.1 and 1.6, respectively) and strong biofilm formation (OD□□□ of 6.1 and 6.7, respectively). Co-cultivation of these two bacterial species resulted in slightly enhanced growth compared to monocultures, both in planktonic (OD□□□ = 2.6) and biofilm-associated states (OD□□□ = 7.5), consistent with cooperative rather than competitive interactions. In the *O. piceae*–*S. oneidensis* consortium, planktonic growth (OD□□□ = 2.5) was comparable to the additive growth of both organisms in monoculture, while biofilm formation exceeded the expected additive value (OD□□□ = 8.3), indicating enhanced surface-associated growth. An even stronger synergistic effect was observed in the *O. piceae–P. putida* consortium, where both planktonic and biofilm growth substantially exceeded the sum of individual cultures (OD□□□ = 2.6 and OD□□□ = 10.5), suggesting a particularly favorable interaction between these two microorganisms. Finally, the three-member consortium exhibited the highest planktonic density (OD□□□ = 3.1) and the second highest biofilm signal (OD□□□ = 9.1) (Fig. 2).

**Figure 2.**
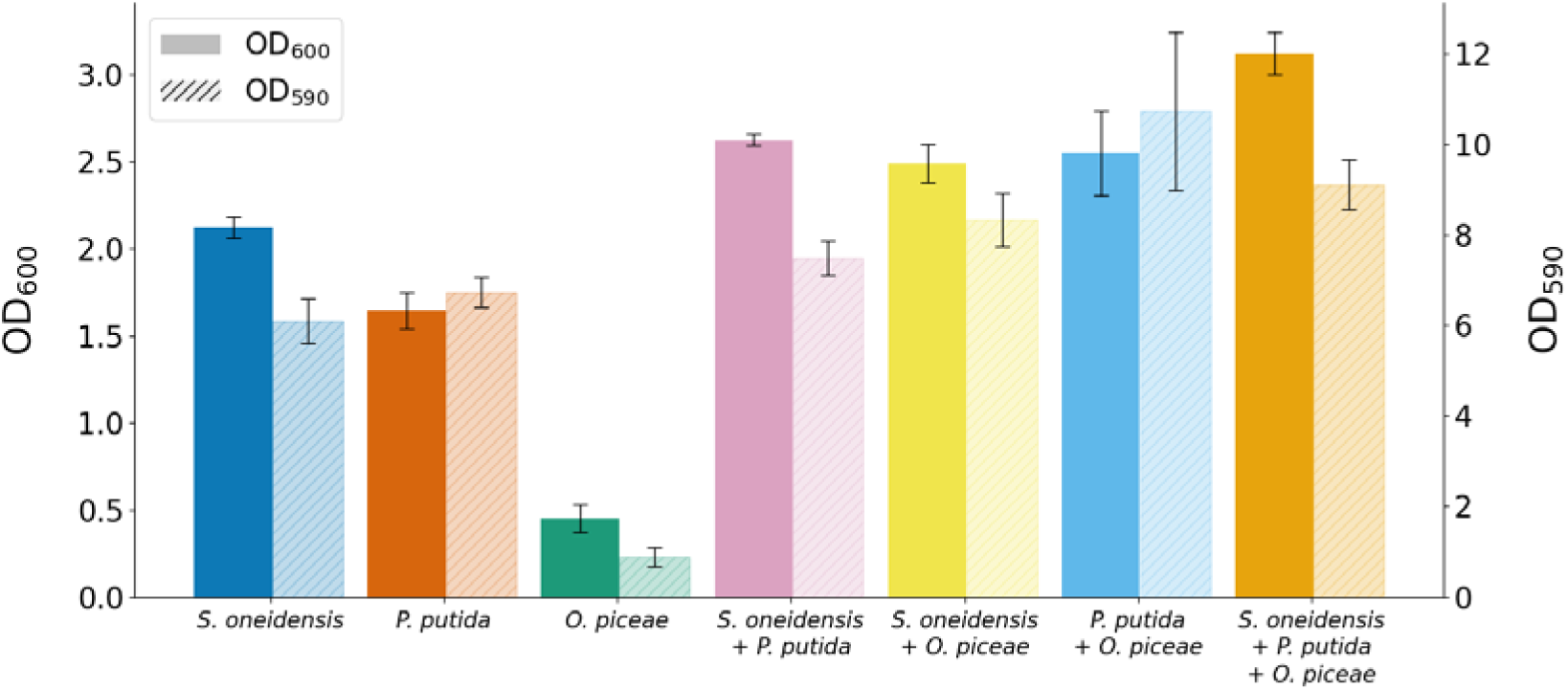
Planktonic growth (OD□□□) and biofilm formation (OD□□□) of the microorganisms used in this study, grown individually or in two- and three-member consortia. Cultures were incubated under static conditions at 30 °C in microtiter plates. Biofilm formation was quantified by crystal violet staining.

Since the ability of microorganisms to adhere to surfaces or form biofilms varies with the properties of the material, the adhesion capacity of the microorganisms under study was examined on carbon veil —the selected electrode material— under non-polarized conditions (i.e., in the absence of a closed circuit, as in an operating MFC) using SEM to characterize the resulting biofilm architecture on the electrode surface. SEM micrographs clearly revealed the adhesion of *P. putida* to the carbon fibers, whereas *S. oneidensis* displayed a comparatively low adhesion capacity (Fig. S1). In contrast, *O. piceae* did not appear to actively adhere to the electrode surface but was mainly observed as hyphae trapped within the carbon fiber network. Notably, the presence of the fungal partner was associated with an increased local presence of *S. oneidensis* on the electrode surface, suggesting that fungal structures may facilitate bacterial retention or attachment. The presence of the triple consortium further promoted the retention of fungal biomass within the electrode structure (Fig.S1). Overall, planktonic growth patterns and SEM observations indicate that the selected microorganisms are compatible and capable of coexisting under non-polarized conditions, with evidence of synergistic interactions in both planktonic and surface-associated states. Differences in apparent growth reflect physiological traits: bacterial biomass can be reliably monitored by OD□□□, whereas *O. piceae*’s filamentous growth and preference for slightly acidic conditions likely limit turbidity measurements without necessarily indicating reduced metabolic activity. These findings suggest that, as hypothesized, the designed anodic medium provides a chemical environment suitable for supporting growth, redox activity, and potential syntrophic interactions, forming a basis for studying multi-species consortia in MFC systems.

### Electrochemical performance of single cultures and microbial consortia

The electrical performance of the MFCs was evaluated by measuring voltage (Figs. 3 and S2), current and power output (Fig. 4) for each microorganism cultivated individually as well as for all possible microbial consortia, using the previously described medium and carbon veil as electrodes. Voltage values at 24 h and 92 h are shown to represent the early stage of operation and the final time point of the experiment in closed circuit configuration, respectively. As expected, the non□ electroactive microorganisms exhibited very poor electrochemical performance under the tested conditions (voltage around 50 mV at 24 and 92 h, current between 0.05 and 0.11 mA and a maximum power of 0.004 mW) whereas *S. oneidensis* exhibited high capacity for extracellular electron transfer to the anode, resulting in an electrical output (104 mV at 24 h and 115 mV at 92 h, 0.25 mA and 0.017 mW). However, cultivation of the electroactive bacteria with *P. putida* generally resulted in reduced electrochemical performance relative to the monoculture of *S. oneidensis* (117 and 102 mV at 24 and 92 h respectively, 0.14 mA and 0.014 mW).

**Figure 3.**
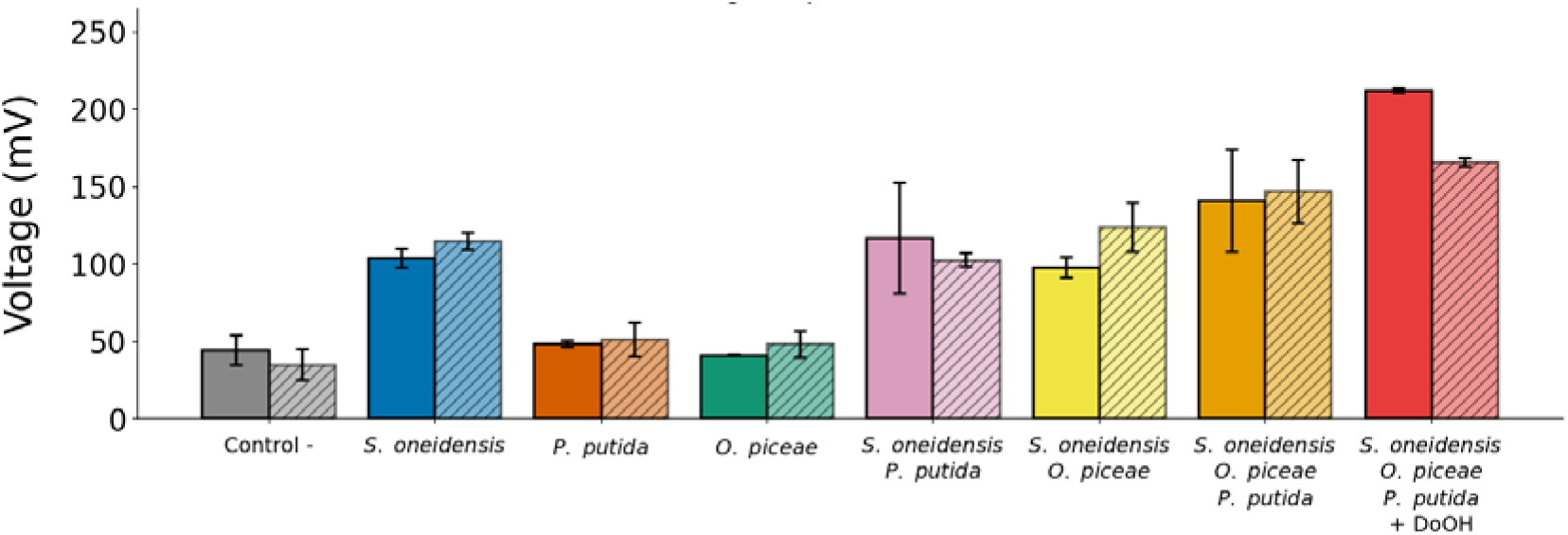
Comparison of voltage output produced by MFCs inoculated with different microorganisms, either as single cultures or consortia, evaluated after 24 h and 92 h of operation. Electricity production of the triple consortium was tested in the absence and presence of the QSM analogue 1-docecanol (DoOH). Solid bars indicate 24 h, and hatched bars indicate 92 h of operation.

**Figure 4.**
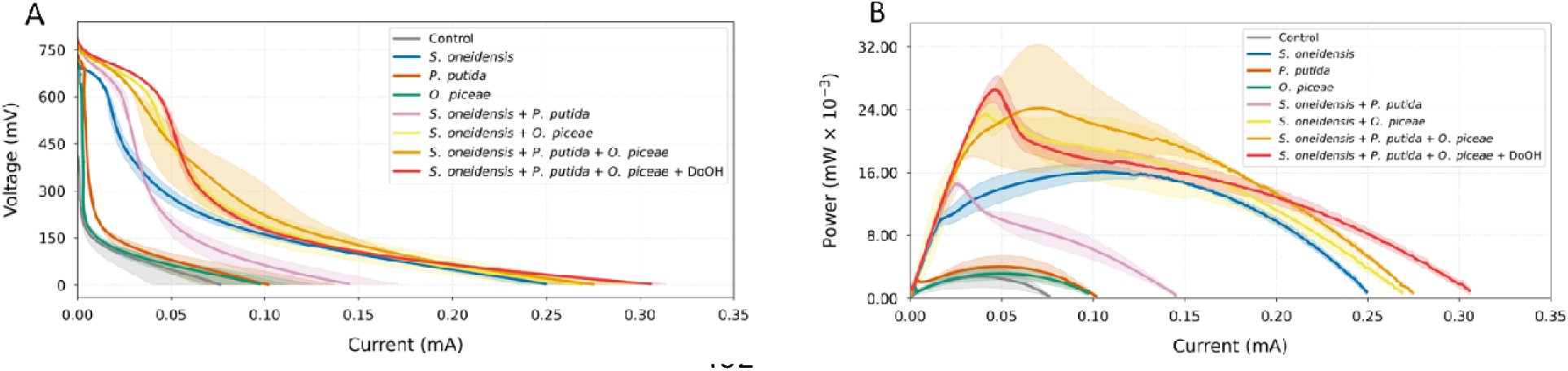
(A) Voltage–current curves and (B) power–current curves obtained for MFCs inoculated with individual microorganisms or with microbial consortia formed by different combinations of the three microorganisms.

Notably, electron transfer was enhanced when *S. oneidensis* was cultivated in consortium with the fungus *O. piceae,* as deduced from the increased values of most electrical parameters compared to the single-species system (98 and 123 mV at 24 and 92 h respectively, 0.27 mA and 0.023 mW). A slightly better electrochemical performance was observed for the triple consortium combining *S. oneidensis, P. putida,* and *O. piceae*. In this configuration, voltage at 24 h was 141 mV and 147 mV after 92 h, current values exceeded 0.27 mA and power output approached 0.024 mW.

Analysis of these results suggests that bioelectrochemical performance is strongly influenced by community structure. Only systems containing *S. oneidensis* generated substantial current, confirming that EET was primarily driven by this electroactive bacterium. The presence of *P. putida* alone reduced electrical output, but co-cultivation with *O. piceae* consistently enhanced current and power density. To further explore strategies for enhancing electrochemical performance, we investigated whether the addition of the QSM analogue 1-dodecanol could improve system output. Based on previous findings (Ruiz et al., 2021), this compound has been reported to modulate interspecies interactions in fungal–bacterial systems by influencing the formation and architecture of mixed biofilms and promoting tighter microbial clustering through effects on community organization. The supplementation of the medium with this QSM resulted in the highest electrochemical performance among all tested conditions, yielding 212 mV at 24 h, 166 mV at 92 h, a maximum current of 0.31 mA and a maximum power output of 0.027 mW. These results suggest that QS-mediated interactions may further stimulate microbial cooperation and extracellular electron transfer, thereby enhancing electricity generation in MFCs.

### Microbial growth dynamics under electricity-generating conditions

After evaluating electricity production, planktonic growth was quantified by measuring optical density in the anolyte (Fig. 5) to assess whether the growth synergies previously observed under oxygen limitation persisted during MFC operation. Surprisingly, *S. oneidensis* population (OD□□□0.7) was notably lower than that observed under non-electrochemical conditions (OD□□□2.1). In contrast, *P. putida*, as expected, displayed robust planktonic growth in the anolyte during MFC operation (OD□□□2.3). As the fungus predominantly grew at the anolyte surface rather than in suspension, the OD□□□ value determined for *O. piceae* was very low (0.38). When microorganisms were cultivated in consortia, the double combinations exhibited higher OD□□□ values than those observed in monocultures (2.9 in the consortium *S. oneidensis* -*P. putida* and 1.1 in the case of *S. oneidensis* - *O. piceae*), a trend already observed in the microplate experiment (Fig. 2). However, the overall planktonic growth in the triple consortium was comparable to that of *P. putida* alone (OD□□□2.2). Interestingly, the addition of 1-dodecanol to the triple consortium resulted in a marked increase in biomass (OD□□□2.9). This value was comparable to those observed for the *S. oneidensis–P. putida* consortium (Fig. 5), suggesting a strong stimulatory effect of this molecule on microbial growth.

**Figure 5.**
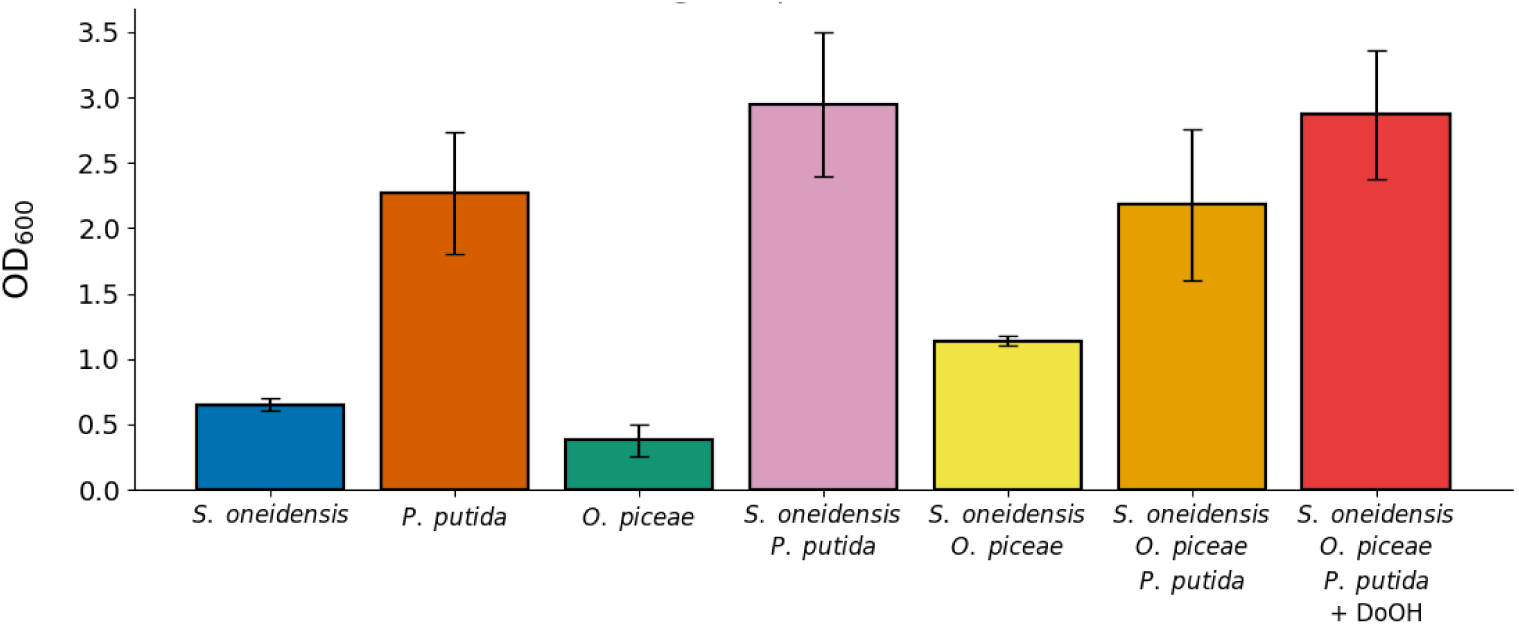
Optical density at 600 nm (ODLJLJLJ) measured in the anolyte of MFCs inoculated with individual microorganisms or with microbial consortia formed by different combinations of the three microorganisms. The effect of supplementation with 1-dodecanol (DoOH) was tested in the triple consortium.

In parallel, the abundance of microorganisms associated with the anode surface was analyzed under the electricity-generating conditions using SEM and CLSM (Fig. 6). The observations revealed a pattern consistent with that previously described under non-polarized conditions. *S. oneidensis* exhibited a relatively low adhesion capacity to the anode surface (Fig. 6A), whereas *P. putida* colonized the electrode much better (Fig. 6B). Since *O. piceae* grows preferentially on the surface of the anolyte, its presence on the anode was quite limited (Fig. 6C). The co-culture of *S. oneidensis* with *O. piceae* did not seem to increase anode adhesion (Fig. 6E). Notably, the overall abundance of bacteria attached to the anode in the triple consortium (Fig. 6F) seemed slightly lower than in *P. putida* or in the *S. oneidensis–P. putida* consortium (Fig. 6D), which exhibited the highest amount of microorganism adhesion, but higher than in *S. oneidensis* monoculture. Furthermore, no noticeable effect of 1LJdodecanol on microbial attachment to the anode was observed (Fig. 6G).

**Figure 6.**
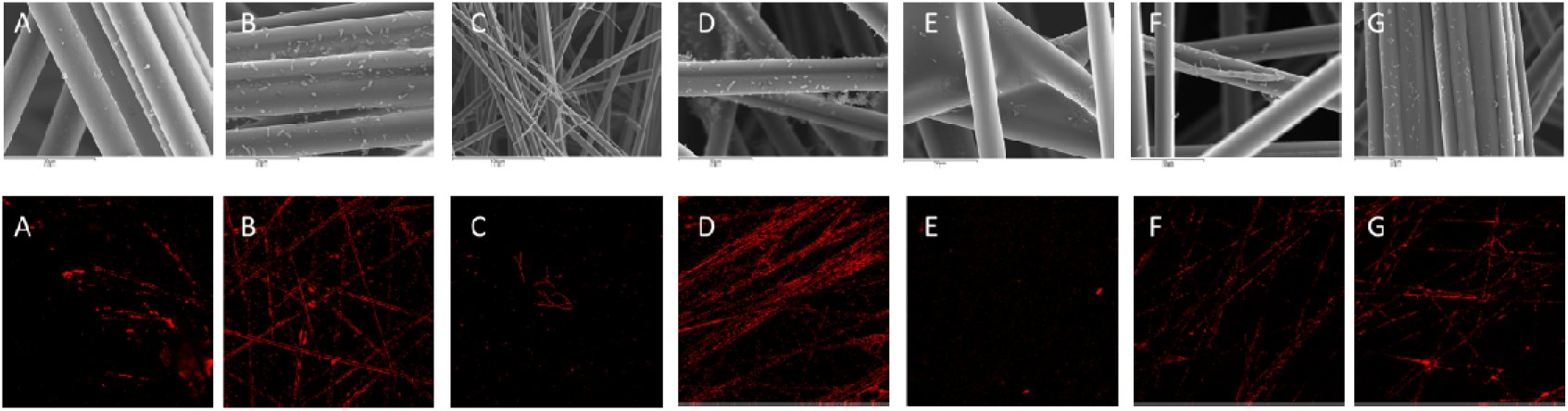
Representative images showing the adhesion of different microorganisms grown individually or in consortia on carbon-veil anodes retrieved from MFCs. SEM micrographs are shown in the upper panels, while CLSM images of the same samples stained with propidium iodide are shown in the lower panels. Panels correspond to individual species: (A) *S. oneidensis*, (B) *P. putida*, and (C) *O. piceae*; and microbial consortia: (D) *S. oneidensis* + *P. putida*, (E) *O. piceae* + *S. oneidensis*, (F) *O. piceae* + *S. oneidensis* + *P. putida*, and (G) the same triple consortium supplemented with 1-dodecanol.

Taken together with the electrical performance data these results support the hypothesis that the fungus acts as a structural scaffold, promoting tighter bacterial clustering, stabilizing mixed biofilms, and favoring metabolite exchange, as previously reported (Ruiz et al., 2021). Hyphal structures trapped within the carbon fiber network appear to compensate for *S. oneidensis* limited adhesion, bringing cells into closer contact with the electrode and indirectly enhancing EET efficiency. Overall, these findings highlight the role of *O. piceae* in architectural and ecological modulation of the microbial community, which contributes to improved electrochemical performance even though the fungus itself does not directly participate in electron transfer.

### Flavin production under electricity-generating conditions

To determine whether the differences in electricity production were driven by *S. oneidensis* adhesion to the anode or by the production of redox mediators by planktonic cells, flavin concentrations in the anolyte were quantified by fluorescence analysis.

As expected, high flavin production was detected in all anolytes containing *S. oneidensis*. The highest riboflavin levels were measured both in the *S. oneidensis* monoculture (444 RFUs) and in consortium with *O. piceae* (420 RFUs), whereas the triple-consortium, without and with 1-dodecanol, showed lower but comparable values (270 and 288 RFUs, respectively) (Fig. 7). Even lower flavin concentrations were detected in the *S. oneidensis–P. putida* consortium (215 RFUs), which also exhibited substantially reduced current output. Riboflavin content in *O. piceae* monocultures did not differ from the microorganism-free control (71 and 61 RFUs respectively), while in *P. putida* it was about twofold higher than the control (139 RFUs) (Fig. 7) which may partially reflect cell lysis, releasing intracellular flavins rather than active secretion.

**Figure 7.**
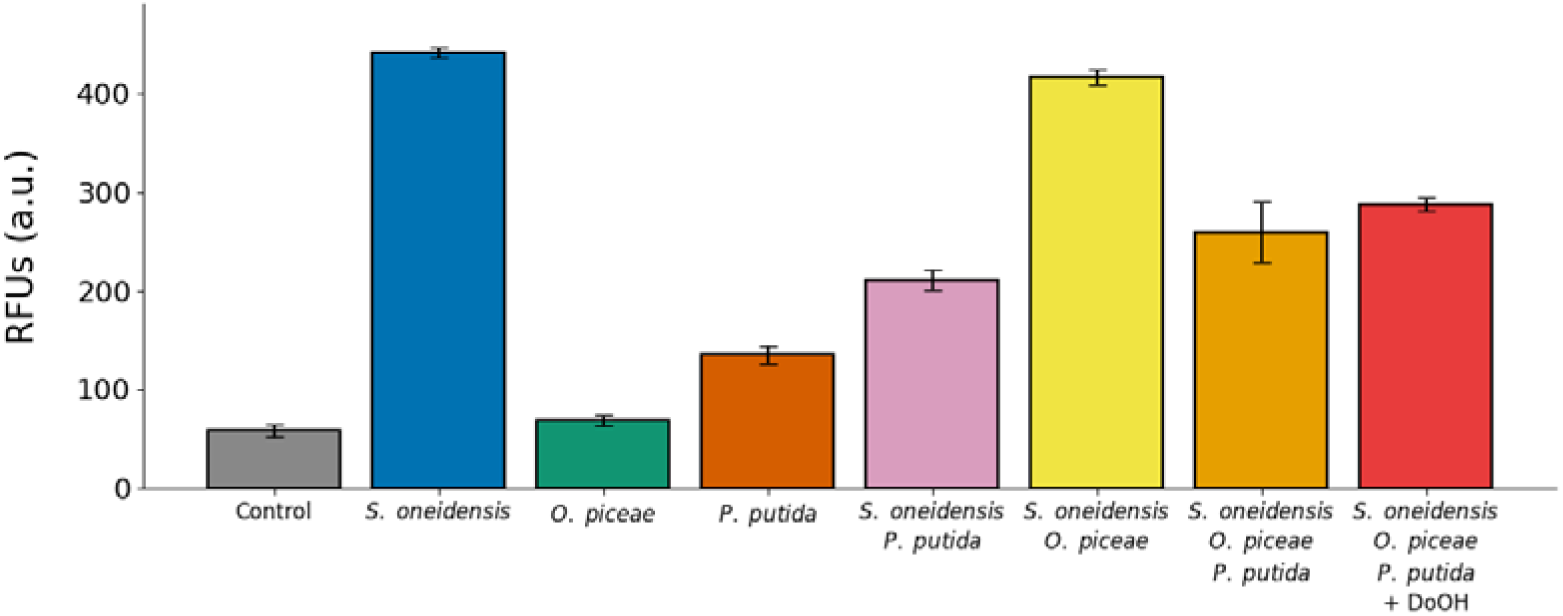
Flavin production, expressed as relative fluorescence units (RFUs), measured in the anolytes of MFCs inoculated with individual microorganisms and different microbial consortia. The effect of supplementation with 1-dodecanol (DoOH) was tested in the triple consortium.

These results indicate that flavin abundance alone does not fully explain electrical performance in the tested MFCs. Instead, the data suggest that the efficiency of EET depends largely on biofilm architecture, the spatial proximity of *S. oneidensis* cells to the electrode. Therefore, bioelectrochemical performance in these consortia appears to arise from the combined effects of mediator availability, microbial adhesion, and community spatial organization, rather than flavin concentration alone.

### Substrate consumption analysis in MFC anolytes

The metabolic activity of the different microbial configurations in the MFCs was investigated by quantifying the consumption of the main components of the anolyte (urea, citric acid and lactic acid) by HPLC and GC–MS (Fig. 8). According to these analyses, lactic acid was almost completely depleted under all tested conditions. Regarding citric acid, the behavior of the different cultures varied, ranging from a lack of assimilation of this compound in *S. oneidensis* monocultures to its total or majority consumption in all those containing *P. putida*. The fungal monoculture used more than half of the available citrate (62%), and in consortium with *S. oneidensis*, consumption was 5.1%. Urea use also differed between microorganisms. *S. oneidensis* was unable to metabolize this nitrogen source, while *O. piceae* and *P. putida* consumed 32% and 10%, respectively. As *S. oneidensis* does not consume urea, its use in all consortia depends on the metabolism of the other members. The values detected in double consortia were very similar to those of the monocultures of *O. piceae* and *P. putida*, and in the triple consortium coincided more or less with the sum of both (38%). In the last case, addition of 1-dodecanol increased urea consumption by 5%, which supports the enhanced growth detected in these conditions (Fig. 5).

**Figure 8.**
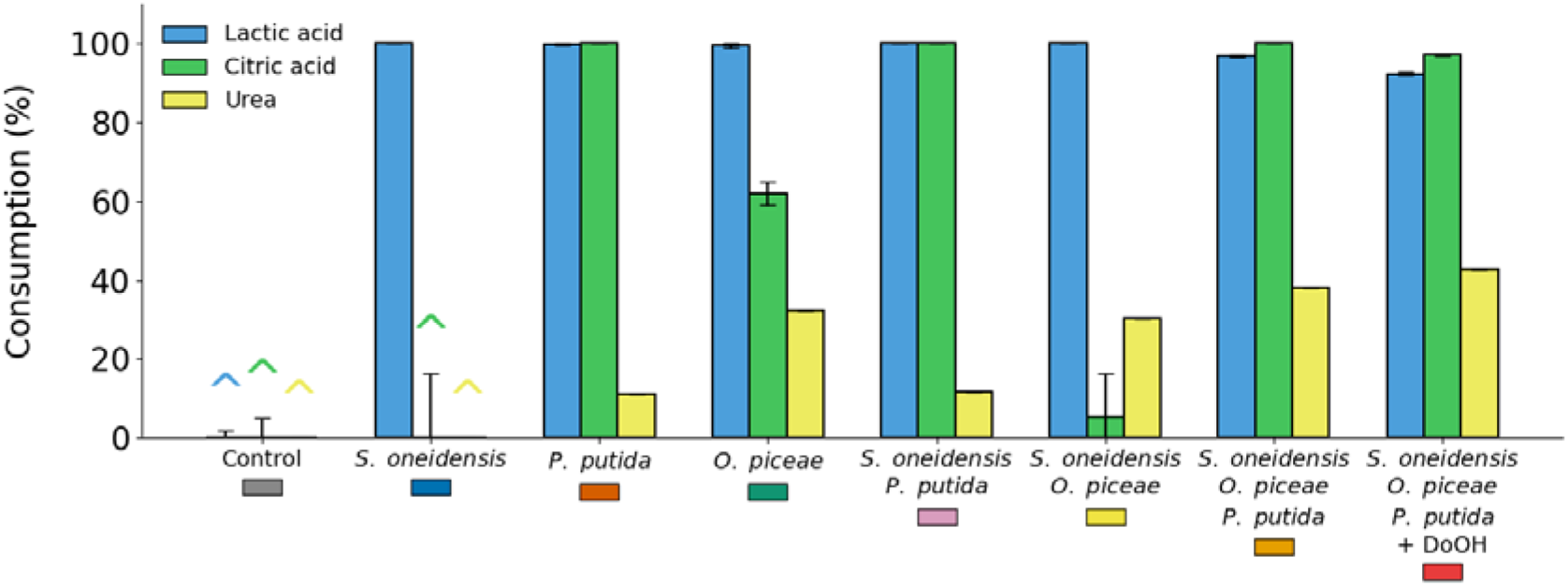
Consumption of lactic acid, citric acid, and urea measured in the anolytes of MFCs inoculated with individual microorganisms or with microbial consortia formed by different combinations of the three microorganisms. The effect of supplementation with 1-dodecanol (DoOH) was tested in the triple consortium.

These metabolic patterns highlight community complementarity in substrate utilization. Lactate was universally consumed, while citrate assimilation was species-dependent and most pronounced in *P. putida*- and *O. piceae*-containing cultures, reflecting their broader metabolic versatility. Urea consumption was primarily associated with nitrogen metabolism rather than carbon oxidation, with the triple consortium showing the highest overall depletion, suggesting cooperative nitrogen turnover. The broader substrate utilization observed in mixed consortia likely contributed to higher biomass formation and improved electrochemical stability, linking effective removal of wastewater components with electricity generation.

### Siderophore production under MFC operating conditions

Siderophores are plant growth–promoting metabolites due to its relevance for improving iron availability in hydroponic cultures. To date, production of such compounds has not been described for either *S. oneidensis* or *O. piceae*, whereas *P. putida* is known to release pyoverdine and pseudobactin, siderophores with natural fluorescence. Thus, the presence of these molecules in the anolyte under MFC operation was assessed by fluorescence analysis, in the presence and absence of EDTA (Fig. 9). In this context, EDTA was added to chelate iron from siderophore–iron complexes, thereby releasing the free siderophores and enhancing the signal. Fluorescence measurements revealed the production of siderophores in all cultures containing *P. putida*. The triple consortium exhibited fluorescence levels comparable to those of the *P. putida* monoculture (486 and 500 RFUs respectively with EDTA). Supplementation with 1-dodecanol led to a marked increase in the fluorescence signal of the triple consortium (977 RFUs with EDTA), which was only surpassed in the *S. oneidensis–P. putida* consortium (1076 RFUs with EDTA). Darker bars represent no EDTA condition while lighter bars indicate the 2mM EDTA condition.

**Figure 9.**
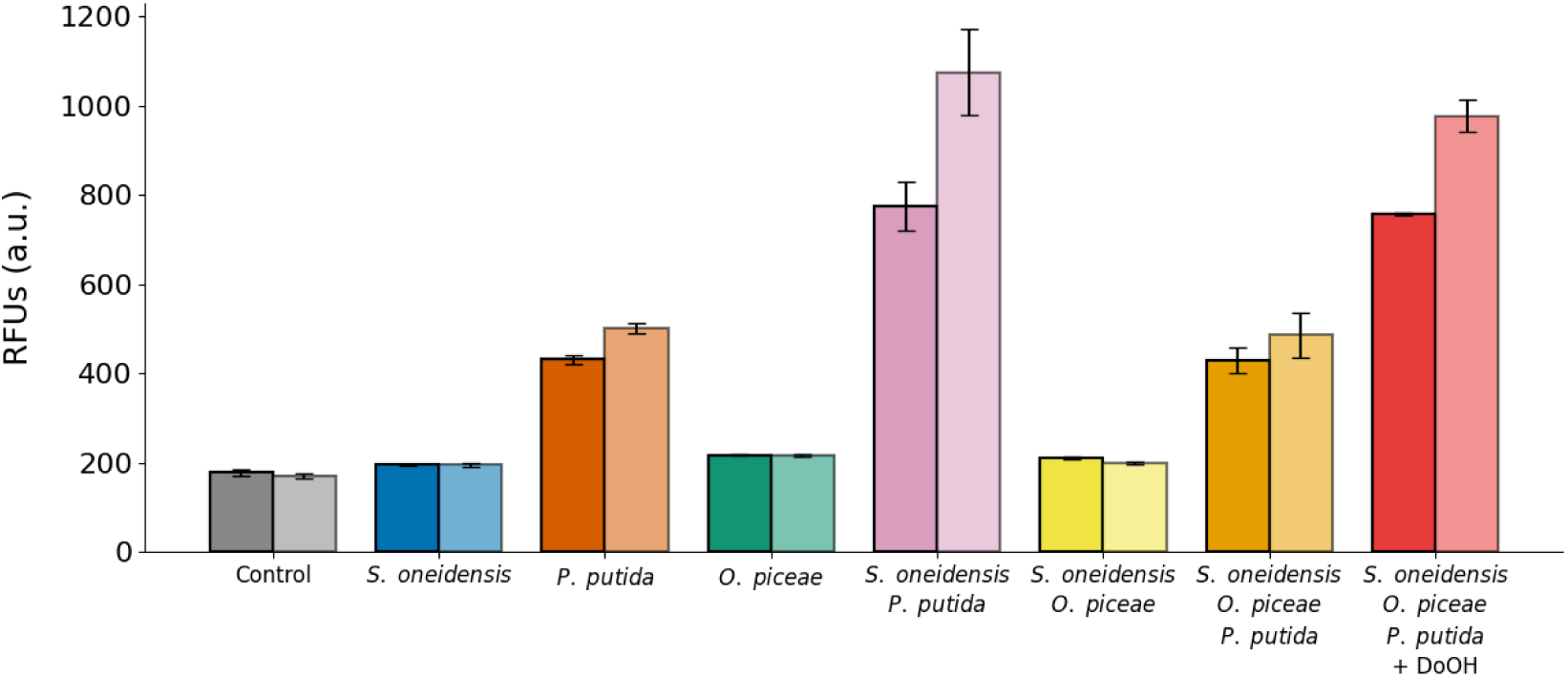
Siderophore production, expressed as relative fluorescence units (RFUs), measured in the anolytes of inoculated with individual microorganisms or with microbial consortia formed by different combinations of the three microorganisms. The effect of supplementation with 1-dodecanol (DoOH) was tested in the triple consortium.

These results confirm that *P. putida* retains an active iron-acquisition metabolism within mixed communities and that siderophore production is maintained, and even enhanced, under certain consortium configurations, likely due to competitive stress for iron when coexisting with other microorganisms, as reported for other species within the same genus (Harrison et al., 2008). The stimulation observed upon addition of 1-dodecanol suggests that QS–related interactions may modulate siderophore production, further supporting cooperative behavior within the community. Altogether, these findings reinforce the concept of the anode as a prosthetic rhizosphere, in which microbial activity not only contributes to bioelectrochemical performance but also generates metabolites with potential relevance for plant nutrition.

## CONCLUSIONS

This study demonstrates that integrating MFCs into a circular wastewater–hydroponic platform enables simultaneous wastewater treatment, electricity generation, and the production of metabolites relevant for plant growth within a single interconnected system. Unlike conventional unidirectional configurations (Nath & Ieropoulos, 2025; Paucar & Sato, 2022; Sato et al., 2023), the bidirectional coupling between the MFC and the hydroponic unit establishes a prosthetic rhizosphere in which microbial metabolism at the anode not only removes organic matter and generates energy, but also releases compounds beneficial for plants, while root exudates in turn sustain anodic microbial activity. Within this framework, a synthetic consortium combining an electroactive bacterium, a metabolically versatile PGPR and a dimorphic fungus displayed cooperative interactions that enhanced substrate utilization and improved bioelectrochemical performance compared with monocultures. Electrical output depended jointly on flavin-mediated electron transfer, electrode colonization, and interspecies interactions, and QS modulation with 1-dodecanol further stimulated system performance.

Taken together, these results suggest that oxygen-dependent trophic interactions are likely established in the system. The fungus may remain near the surface, metabolizing substrates and extending hyphae toward the electrode, while *P. putida*, as an obligate aerobe, persists mainly in planktonic form, producing siderophores and consuming residual oxygen. Both microorganisms likely contribute indirectly to extracellular electron transfer by *S. oneidensis* through nutrient recycling, redox modulation, and metabolic support. Supplementation with 1□ dodecanol may enhance these interactions by modulating interspecies communication, promoting tighter cell clustering, and increasing overall metabolic cooperation.

Importantly, this study was not designed to maximize absolute power density but to evaluate microbial consortia within a multifunctional circular wastewater–hydroponic framework. Within this context, the triple mixed consortium emerged as the most balanced configuration, exhibiting superior substrate removal, metabolite production, and electricity generation. To our knowledge, this represents the first circularly integrated MFC–hydroponic platform recreating a prosthetic rhizosphere and incorporating a dimorphic fungus, highlighting the potential of rationally designed multispecies communities to enhance both ecological functionality and bioelectrochemical performance.

## ACKNOWLEDGEMENTS

This work has been funded by the European project Mi-Hy (PATHFINDERCHALLENGES-01 EIC 101114746). The authors acknowledge the support toward the publication fee by the CSIC Open Access Publication Support Initiative through its Unit of Information Resources for Research (URICI). DGG acknowledges financial support from Fundación Ramón Areces for his PhD project.

