## Supplementary Material for "Integrating Fungal–Bacterial Synergy to Enhance Circular MFC–Hydroponic Performance"

**Table S1. Formulation of the anodic medium employed in this study**

| **Component** | **Concentration** |
| --- | --- |
| Ca(NO_3_)_2_ · 3H_2_O | 5.04 mM |
| NH_4_NO_3_ | 1.08 mM |
| KNO_3_ | 9.9 mM |
| EDTA iron(III) sodium salt | 12.24 µM |
| MnSO_4_ · 7H_2_O | 0.45 µM |
| H_3_BO_3_ | 3.375 µM |
| NaMoO_4_ · 2H_2_O | 0.0567 µM |
| ZnSO_4_ · 7H_2_O | 0.45 µM |
| CuSO_4_ · 5H_2_O | 0.0837 µM |
| Citric acid | 0.9 mM |
| Lactic acid | 0.495 mM |
| NaCl | 40.5 mM |
| NH_4_Cl | 11.25 mM |
| RPMI 1640 aa solution | 9 mL/L |
| L-glutamine | 0.9 mM |
| Urea | 76.5 mM |
| Uric acid | 0.18 mM |
| Creatinine | 3.15 mM |
| CaCl_2_ · 2H_2_O | 0.112 mM |
| MgSO_4_ · 7H_2_O | 3.17 mM |
| Na2SO_4_ · 10H_2_O | 4.5 mM |
| NaHCO_3_ | 11.25 mM |
| NaNO_3_ | 2.7 mM |
| FeSO_4_ · 7H_2_O | 2.25 µM |
| KH_2_PO_4_ | 3.08 mM |
| K_2_HPO_4_ | 0.81 mM |
| Peptone | 0.45 % (w/v) |
| Yeast extract | 0.225 % (w/v) |
| Cellulose | 0.00625 % (w/v) |
| Pectin | 0.0025 % (w/v) |
| Xylan | 0.00125 % (w/v) |
| Ferulic acid | 0.00148 µM |
| Pipecolic acid | 0.00154 µM |
| Allantoin | 0.00126 µM |
| Histamine | 0.00179 µM |

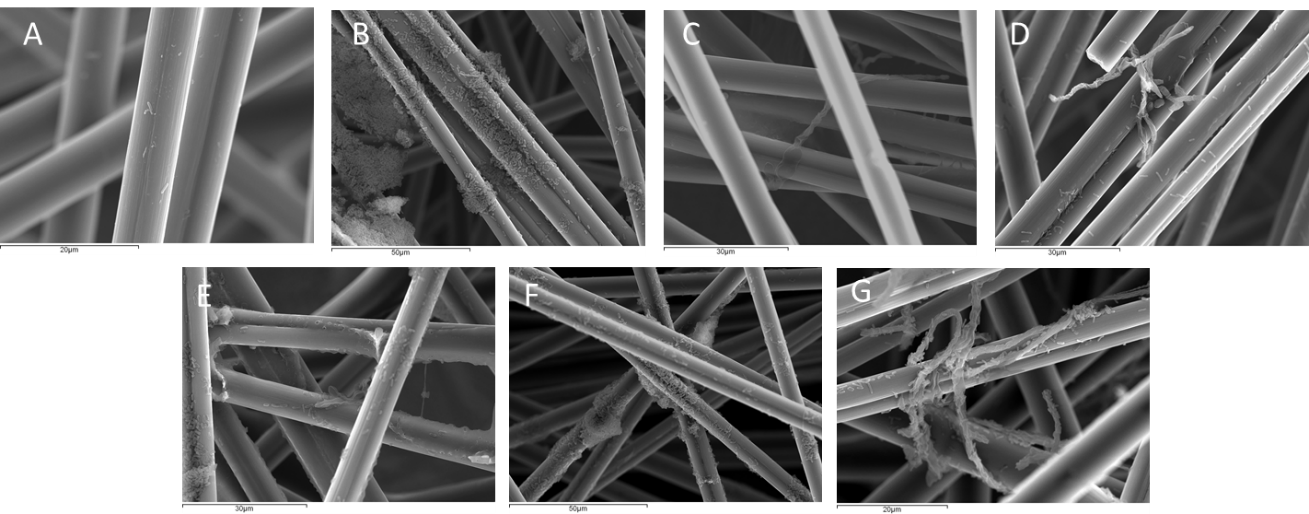

**Figure S1.** Representative SEM images showing the adhesion of the different microorganisms grown individually or in consortia on carbon‑veil anode material under non‑poised conditions. Panels correspond to individual species: (A*) S. oneidensis*, (B*) P. putida*, (C) *O. piceae*; and microbial consortia: (D) *O. piceae* + *S. oneidensis*, (E) *O. piceae + P. putida*, (F) *S. oneidensis + P. putida*, and (G) *O. piceae + S. oneidensis + P. putida*.

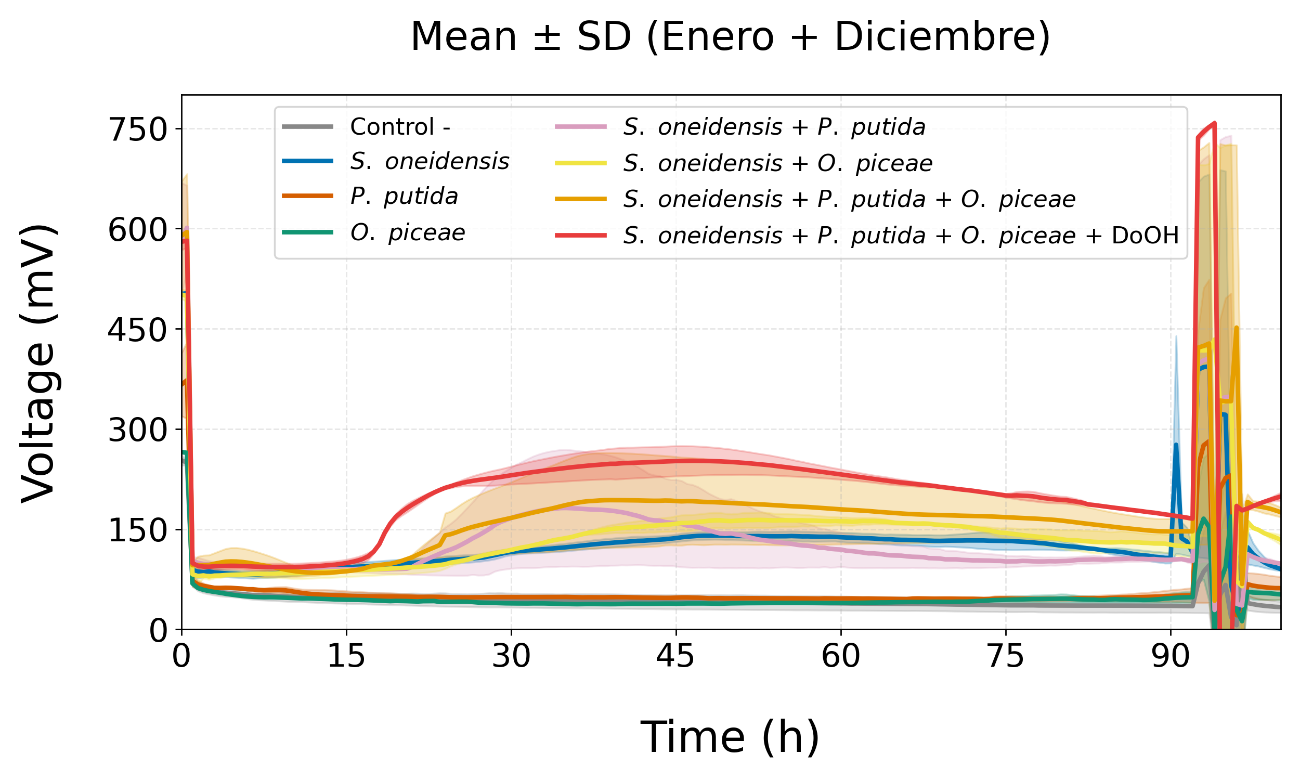

**Figure S2**. Comparison of voltage output produced in 96 h by microbial fuel cells (MFCs) inoculated with individual microorganisms or with microbial consortia formed by different combinations of the three microorganisms. The effect of supplementation with 1-dodecanol (DoOH) was tested in the triple consortium.
